# The cell walls of different *Chara* species (Charophyceae) are characterized by branched galactans rich in 3-*O*-methylgalactose and absence of arabinogalactan-proteins

**DOI:** 10.1101/2023.05.22.541140

**Authors:** Lukas Pfeifer, Kim-Kristine Mueller, Jon Utermöhlen, Felicitas Erdt, Jean Bastian Just Zehge, Hendrik Schubert, Birgit Classen

**Author notes:** Author for correspondence: Birgit Classen, Pharmaceutical Institute, Department of Pharmaceutical Biology, Christian-Albrechts-University of Kiel, Gutenbergstr. 76, 24118 Kiel, Germany. Authors contributed equally to the publication (Lukas Pfeifer, Kim-Kristine Mueller).

## Abstract

Streptophyte algae are the closest relatives of land plants and their latest common ancestor performed the most drastic adaptation that happened in plant evolution around 500 million years ago: the conquest of land. Beside other adaptations, this step required changes in cell wall composition. Today knowledge on cell walls of streptophyte algae and especially presence of arabinogalactan-proteins (AGPs), which are important signaling molecules of all land plants, is limited. To get deeper insights in cell walls of streptophyte algae, especially of the Charophyceae, we performed sequential cell wall extractions of four *Chara* species. The three species *Chara globularis*, *Chara subspinosa* and *Chara tomentosa* revealed comparable cell wall compositions with pectins, xylans and xyloglucans, whereas *Chara aspera* was outstanding with higher amounts of uronic acids in the pectic fractions and lack of reactivity with antibodies binding to xylan- and xyloglucan epitopes. Search for AGPs in the four *Chara* species and also *Nitellopsis obtusa* revealed presence of galactans with pyranosidic galactose in 1,3-, 1,6- and 1,3,6-linkage, which are typical galactan motifs of land plant AGPs. A unique feature of these branched galactans were high portions of 3-*O*-methylgalactose. Only *Nitellopsis* contained substantial amounts of Ara. Bioinformatic search for prolyl-4-hydroxylase necessary for biosynthesis of AGPs revealed one possible functional sequence in the genome of *Chara braunii*, but no hydroxyproline could be detected in the four *Chara* species and *Nitellopsis obtusa*. We conclude that AGPs typical for land plants are absent at least in these members of the Charophyceae.

## INTRODUCTION

About more than 500 million years ago (mya) one of the most significant events for life on Earth occurred: the terrestrialization of land (Domozych & Bagdan, 2022). Land plants (bryophytes, lycophytes, ferns, gymnosperms, and angiosperms) are descended from a common ancestor belonging to the streptophyte algae (Bowman, 2022), which are divided into two grades: the lower-branching KCM-grade (Klebsormidiophyceae, Chlorokybophyceae and Mesostigmatophyceae) and the higher-branching ZCC-grade (Zygnematophyceae, Coleochaetophyceae and Charophyceae; de Vries & Archibald, 2018). The genus *Chara* is an important constituent of natural aquatic ecosystems, commonly beneficial to lakes and ponds by providing food and cover to wildlife (DiTomaso & Kyser *et al*., 2013). Due to the morphological complexity of the Charophyceae, it was first assumed that these streptophyte algae were the closest ancestors of extant land plants (Domozych & Bagdan, 2022). However, recent phylogenomic studies have shown that land plants evolved from streptophyte algae most closely related to extant Zygnematophyceae (Domozych & Bagdan, 2022; Leebens-Mack *et al*., 2019; Wickett *et al*., 2014), which comprise a five-order system including Desmidiales, Spirogyrales, Zygnematales, Serritaeniales and Spirogloeales (Hess *et al*., 2022).

To cope with life on land, the terrestrial algal species had to adapt to enormous changes in the environment, such as UV-radiation and drought. This also meant that the cell wall needed to adapt by changing its composition (Harholt *et al*., 2016). The plant cell wall is generally a highly dynamic system consisting mainly of polysaccharides as well as of cell wall proteins. It has many essential functions such as mechanical stability and protection against abiotic and biotic stressors (Silva *et al*., 2020). The main polysaccharides of spermatophyte cell walls include cellulose, pectins, hemicelluloses and, in vascular plants, lignin. The hemicelluloses are cross-linked in a highly branched network of cellulose microfibrils, which are embedded in a matrix of pectins (Sørensen *et al*., 2011). The cell wall proteins consist mainly of hydroxyproline-rich glycoproteins (HRGPs), like arabinogalactan-proteins (AGPs), proline rich glycoproteins (PRPs) and extensins (EXT; Johnson *et al*., 2018). The three HRGP-classes differ mainly in their carbohydrate moiety. While this part is about 90% in AGPs, it is about 50% in EXT and in PRPs, the glycan part is only marginally glycosylated (Johnson *et al*., 2018).

The AG moiety of AGPs consists of arabinogalactans type II (AG II) with a (1→3)-β-D-galactan core structure carrying (1→6)-β-D-galactan side chains at position O-6. These side chains are decorated with α-L-Ara*f* and other monosaccharides, like L-Rha, L-Fuc or D-GlcA*p* (Kitazawa *et al*., 2013). The AG IIs are covalently linked *via* hydroxyproline (Hyp) to the protein moiety. The hydroxylation of Pro to Hyp is post-translationally catalyzed by prolyl 4-hydroxylase (P4H; Seifert *et al*., 2021).

Additionally, a glycosylphosphatidylinositol (GPI)-anchor can be attached to the C-terminal sequence, which connects the AGP to the plasma membrane (Silva *et al*., 2020). Even if it is not yet fully explained how the signal transmission of AGPs takes place, this localization nevertheless facilitates this process. AGPs of seed plants show many and diverse functions, which include cell growth, pattern formation, salt and drought tolerance, cell-cell communication or adhesiveness (for review, see Ma *et al*., 2018; Mareri *et al*., 2018; Leszczuk *et al*., 2023; Seifert & Roberts, 2007). These functions make AGPs ideal candidates for molecules, which were essential for life on land and may have evolved in the common ancestor of land plants. In ferns and bryophytes, the general AGP structure known from seed plants is present, but shows unique variations in the carbohydrate part, e.g. the presence of 3-*O*-methylrhamnose as terminal monosaccharide (Bartels *et al*., 2017; Bartels & Classen, 2017; Fu *et al*., 2007; Mueller *et al*., 2023). Whether AGPs are present in algae is still questionable. Based on bioinformatics screenings of transcriptomes, protein backbones of AGPs have been proven for brown, red and green algae (Johnson *et al*., 2017). Furthermore, AGP glycan structures have been detected by monoclonal antibodies in brown algae (Hervé *et al*. 2016; Raimundo *et al*., 2016) and green algae including some streptophyte algae (Eder *et al*., 2008; Estevez *et al*., 2008; Estevez *et al*., 2009; Domozych *et al*., 2009; Palacio-López *et al*., 2019; Permann *et al*., 2021a, 2021b; Přerovská *et al*., 2021; Ruiz-May *et al*., 2018; Sørensen *et al*., 2011), but it remains open whether these are linked to AGP protein backbones.

To the best of our knowledge, no AGP similar to a land plant AGP has been isolated from a streptophyte algae to date. Closely related rhamnogalactan-proteins (RGPs) are present in cell walls of *Spirogyra pratensis*, a member of the Zygnematophyceae. These RGPs reveal typical AGP features like a Hyp-rich protein moiety covalently bound to 1,3-, 1,6- and 1,3,6-linked galactans and precipitation with Yariv’s reagent. Interestingly, there is nearly complete replacement of arabinose by rhamnose at the periphery of RGPs, leading to a less hydrophilic surface maybe connected with different functions in the freshwater habitat (Pfeifer *et al*., 2022). To continue the search for evolutionary roots of AGPs or AGP-like molecules in streptophyte algae, we searched for these glycoproteins in four *Chara* species (Charophyceae). As hydroxylation of proline is the basis for O-glycosidic linkage between protein- and carbohydrate moiety in AGPs, we also performed bioinformatic search for prolyl-4-hydroxylases. The work is accomplished by sequential extraction of *Chara* cell walls to get insights into general cell wall polysaccharide composition of this genus. The results broaden our general understanding of cell walls of the higher branching streptophyte algae and further enlighten the process of cell wall evolution from algae to land plants.

## MATERIAL AND METHODS

### Plant material and sampling

The streptophyte algae *Chara globularis* THUILL., *Chara subspinosa* RUPR. and *Chara tomentosa* L. were collected in June 2018 in the lake “Lützlower See” in Uckermark (Germany). *Chara aspera* WILLD. was collected in July 2021 in the lake in Lalendorf (Germany). All samples were cleaned with water and freeze-dried (Christ Alpha 1-4 LSC, Martin Christ Gefriertrocknungsanlagen GmbH, Osterode, Germany). Preliminary experiments were carried out with *C. subspinosa* from the Lunzer See and *C. tomentosa* and *C. globularis* from the Neusiedler See. The Austrian samples were collected and identified by Prof. Dr. Michael Schagerl and Barbara Mähnert from the Department for Limnology and Oceanography of the University of Wien.

### Isolation of cell wall fractions

The freeze-dried and ground plant material was two times pre-extracted (2 h and 21 h) with 70% acetone solution (*V/V*). After drying, the plant residue was extracted with double-destilled water (ddH_2_O) for 21 h under constant stirring (SM 2484, Edmund Bühler GmbH, Bodelshausen, Germany) at 4°C in a 1:10 (*w/V*) ratio. The aqueous extract was separated through a tincture press (HAFICO HP 2 H, Fischer Tinkturenpressen GmbH, Mönchengladbach, Germany) and the insoluble plant pellet was freeze-dried for further extractions (see below).

After heating the aqueous extract in a water bath (GFL 1002, LAUDA-GFL Gesellschaft für Labortechnik mbH, Burgwedel, Germany) at 90 – 95°C for ten minutes, the denatured proteins were removed by centrifugation (4,122 *g*, 20 min, 4°C, Haraeus Multifuge X3R, Thermo Scientific Waltham, USA). To precipitate polysaccharides and AGPs, the aqueous extract was first evaporated in a rotary evaporator (40°C, 0,010 mbar; Laborota 4000, Heidolph Instruments GmbH & CO. KG, Schwabach, Germany) to approximately one-tenth of its volume and subsequently added to cooled ethanol resulting in a final concentration of 80% (*V/V*). After incubation overnight at 4°C, the precipitate was isolated by centrifugation (4°C, 19,000 *g*, 20 min) and finally freeze-dried (AE).

To remove starch and to cleave homogalacturonan parts, approximately 100 mg of AE were treated with α-amylase + amyloglucosidase (α-amylase from *Aspergillus oryzae*, EC number 3.2.1.1., 15 U mg^-1^ polysaccharide; amyloglucosidase from *Aspergillus niger*, EC number 3.2.1.3., 7 U mg^-1^ polysaccharide; both from Sigma-Aldrich Chemie GmbH, Taufkirchen, Germany; in 50 mmol L^-1^ acetate buffer at pH 5.2, 50°C for 20 h) and subsequently with pectinase (Sigma-Aldrich Chemie GmbH, Taufkirchen, Germany, from *Aspergillus niger*, EC number 3.2.1.15., 1µl mg^-1^, for 5h, in 50 mmol L^-1^ acetate buffer at pH 5.2, 37.5°C). Afterwards, the enzymes were removed by heating in a boiling water bath for 10 min with following centrifugation at 19,000 *g* for 20 min. The samples were dialyzed against demineralized water (4°C; 4d; MWCO 12 – 14 kDa, Visking®, Medicell International Ltd, London, UK) and freeze-dried (AE-AP).

According to a modified method of O’Rourke *et al*. (2015) and Raimundo *et al*. (2016), the insoluble plant pellet after water extraction (see above) was used to gain four further polysaccharide fractions. At first, the pellet was subsequentially extracted with ammonium oxalate ((NH_4_)_2_C_2_O_4_, 0.2 mol L^-1^), hydrochloric acid (HCl, 0.01 mol L^-1^), sodium carbonate (Na_2_CO_3_, 3% (*w/V*)) and potassium hydroxide (KOH, 2 mol L^-1^), each in a ratio of 1:100 (*w/V*) at 70 °C under constant stirring for 21 h (RET basic, ETS D5, IKA Labortechnik, Staufen, Germany). The pellet and the supernatant were separated after each extraction by centrifugation (4°C, 19,000 *g*, 20 min, Heraeus Multifuge X3, Thermo Fisher Scientific Corp., Waltham, MA, USA).

To precipitate the Na_2_CO_3_ fraction, the supernatant was added to acetone in a 1:4 (*V/V*) ratio. After incubation overnight at 4°C, the precipitated material was re-dissolved in ddH_2_O in a 1:100 (*w/V*) ratio. The KOH extract was neutralized (761 Calimatic, Knick Elektronische Messgeräte GmbH & Co. KG, Berlin, Germany). Finally, the four fractions were dialyzed against demineralized water (4°C, MWCO 12 – 14 kDa (Visking®), 4 d) and freeze-dried.

### Analysis of monosaccharides

The neutral monosaccharide composition was determined by the method of Blakeney *et al*. (1983) with slight modifications (see Mueller *et al*., 2023).

Gas chromatography (GC) with flame ionization detection (FID) and mass spectrometry detection (MSD) was used to identify and quantify the neutral monosaccharides: GC + FID: 7890B; Agilent Technologies, USA; MS: 5977B MSD; Agilent Technologies, USA; column: Optima-225; Macherey-Nagel, Germany; 25 m, 250 μm, 0.25 μm; helium flow rate: 1 ml min^-1^; split ratio 30:1. A temperature gradient was performed to achieve peak separation (initial temperature 200°C, subsequent holding time of 3 min; final temperature 243°C with a gradient of 2°C min^-1^). The methylated monosaccharide 3-*O*-MeRha was identified by retention time and mass spectrum (see Happ & Classen, 2019), 3-*O*-MeFuc was determined using the mass spectrum and the deviating retention time to 3-*O*-MeRha. 3-*O*-MeGal was also quantified by mass spectrum, but additionally the validation method of Pfeifer *et al*. (2020) was used to distinguish 3-*O*-MeGal from 4-*O*-MeGal.

The uronic acid (UA) compounds were determined photometrically at 525 nm (UVmini-1240, Shimadzu AG, Kyoto, Japan) according to the method of Blumenkrantz & Asboe-Hansen (1973), for modifications see Mueller *et al*. (2023).

### Reduction of uronic acids

A modified method of that of Taylor and Conrad (1972) was used to perform a carboxy-reduction of the uronic acids of the AE_AP fractions. 20 – 30 mg AE_AP were dissolved in 20 mL ddH2O and 216 mg of N-cyclohexyl-N’-[2-(N-methylmorpholino)-ethyl]-carbodiimide-4-toluolsulfonate was added slowly under constant stirring. An autotitrator (Metrohm 719 S-Titrino, Deutsche METHROM GmbH & Co. KG, Filderstadt, Germany) adjusted the pH to 4.75 with 0.01 M HCl for 2 h. First, one to two drops of 1-octanol were added to avoid strong foaming and then sodium borodeuteride solutions in increasing concentrations (4.0 mL of 1 mol L^-1^; 5 mL of 2 mol L^-1^; 5 mL of 4 mol L^-1^) were admitted to reduce the uronic acids. The autotitrator adjusted the pH to 7.00 with 2 M HCl for another 2 h. After pH setting to 6.5 with glacial acetic acid, the solutions were dialysed for three days at 4 °C against demineralized water (MWCO 12 – 14 kDa, Visking®) and freeze-dried.

### Structural characterization of water soluble polysaccharides

Following the method of Harris *et al*. (1984; see Mueller *et al*., 2023 for modifications), structural characterization of the polysaccharides was performed. First, potassium methylsulfinyl carbanion (KCA) and iodomethane-d_3_ (IM-d_3_) were added to the samples in a defined scheme to methylate the samples (1. 100 μl KCA for 10 min; 2. 80 μl IM-d_3_ for 5 min; 3. 200 μl KCA for 30 min; 4. 150 μl IM-d_3_ for 60 min). After subsequent hydrolysis, reduction, and acetylation, the samples were modified to permethylated alditol acetates. These were identified by GLC-mass spectroscopy (instrumentation see above: “Analysis of monosaccharides”; column: Optima-1701, 25 m, 250 µm, 0.25 µm; helium flow rate: 1 mL min^-1^; initial temperature: 170°C; hold time 2 min; rate 1°C min^-1^ until 210°C was achieved; rate: 30°C min^-1^ until 250°C was achieved; final hold time 10 min) by their retention times and comparison to a PMAA library established in the working group.

### Determination of hydroxyproline content

For hydroxyproline (Hyp) quantification, the methodology of Stegemann & Stalder (1967) was used and modified (see Mueller *et al*., 2023). The Hyp content was measured photometrically at 558 nm and quantified *via* a linear regression analysis (standard: 4-hydroxy-L-proline).

### Indirect enzyme-linked immunosorbent assay (ELISA)

After coating of the 96-well plates (Nunc-Immuno® Plates, Thermo Scientific, Waltham, USA) with 100 µl per well of the sample in triplicate in different concentrations in ddH2O (12.5 µg mL^-1^ for AmOx and KOH; 12.5 µg mL^-1^, 25 µg mL^-1^ 50 µg mL^-1^ for AE), these were incubated at 37.5°C for 3 days. The plates were washed three times with 100 µl phosphate buffered saline (PBS)-T (pH 7.4, 0.05% Tween® 20) per well and blocked with 200 µl of BSA (bovine serum albumin, 1% (*w/V*)) in PBS per well. After incubation at 37.5°C for 1 h, the plates were washed again three times. Following the addition of 100 µl of primary antibody solution (INRA-RU2 (1:200 (*V/V*), JIM13 (1:40 (*V/V*)), KM1, LM2, LM6, LM10, LM15, LM19 (1:20 (*V/V*)) per well in PBS 7.4 dilution, the plates were incubated again (1 h at 37.5°C). After washing again three times, the procedure was repeated for the secondary antibody (anti-rat-IgG (others) or anti-mouse-IgG (INRA-RU2; KM1), both produced in goat, conjugated with alkaline phosphatase, Sigma-Aldrich Chemie GmbH, Taufkirchen, Germany) in a ratio of 1:500 (*V/V*) in PBS 7.4. Subsequently, the substrate *p*-nitro-phenylphosphate was added (100 µl well^-1^). The plates were incubated at room temperature in the dark and the absorbance was determined at 405 nm with a plate reader (Tecan Spectra Thermo, Männedorf, CH) after 40 min. For demonstration, the absorbance of the control was set 0 and the highest signal in the dataset was set 1.0. Epitopes of the antibodies and key references are listed in Table S1.

### Gel diffusion assay

For the gel diffusion assay, an agarose gel (Tris-HCl, 10 mmol L^-1^; CaCl_2_, 1 mmol L^-1^; NaCl, 0.9% *w/V*; agarose, 1% *w/V*) was used, in which several cavities were stamped in. The cavities in the first and third row were filled with dilutions (100 mg mL^-1^) of the samples. In the second line, the cavities left and right were filled with the positive control, AGP from *Echinacea purpurea* (10 mg mL^-1^), the middle cavities with the βGlcY solution (1 mg mL^-1^). The assay was incubated for 22 h in the dark. If AGPs are present in the samples, a red precipitation line appears, as in the positive control.

### Bioinformatic search for prolyl-4-hydroxylase (P4H)

Candidate sequences for P4Hs were obtained from the translated proteome of *Chara braunii* (Nishiyama *et al*., 2018) under the use of the R package rBLAST (Hahsler & Nagar, 2019) with the *A. thaliana* P4Hs as query sequences. An E-value of 1e^-7^ was used and the resulting candidates were aligned with functionally described P4H sequences (Hieta & Myllyharju, 2002; Keskiaho *et al*., 2007; Vlad *et al*., 2010; Velasquez *et al*., 2014; Yuasa *et al*., 2005) by the use of MAFFT in L-INS-i mode (Katoh & Standley, 2013). Occurrence of relevant amino acids was evaluated in comparison to *Chlamydomonas reinhardtii* P4H1. Afterwards some sequences were modelled with the SWISS-MODEL webserver (Waterhouse *et al*., 2018) with default settings on the JIG2/A crystal structure of *C. reinhardtii* P4H (Koski *et al*., 2007).

## RESULTS

### Sequential extraction of polysaccharide fractions from *Chara* sp. cell walls

We examined four *Chara* species to investigate their cell wall compositions. Sequential extraction of the plant material with water (AE), ammonium-oxalate (AmOx; (NH_4_)_2_C_2_O_4_) and hydrochloric acid (HCl) as well as sodium carbonate (Na_2_CO_3_) and potassium hydroxide (KOH) resulted in a water-soluble, two pectic and two hemicellulosic fractions. The yields of the different fractions varied between 0.2% and 21.2% of the dry plant material (Table S2). The high yield of the AmOx fraction of *Chara aspera* (21.2%) was outstanding and indicated huge amounts of pectins in this species.

### Pectic fractions of *Chara* sp. cell walls

The monosaccharide compositions of the pectic fractions are shown in Tables 1, 2a and 2b and Figure 1. As expected, the two pectic fractions showed the highest amounts of uronic acids. In the AmOx fractions, their content reached 33% to 60% of the dry fraction material and decreased in the HCl fractions of all investigated *Chara* spp. (Table 1). In contrast to the other *Chara* species, the uronic acid content in *C. aspera* was extraordinary high with over 60% in the AmOx and almost 40% in the HCl fraction. The dominant neutral monosaccharide of both fractions was Glc with 25% to 66% (Tables 2a and 2b). In the HCl fractions, the content was approximately doubled compared to the AmOx fractions. It cańt be excluded that at least part of Glc derived from coextracted starch. Other monosaccharides like Rha, Gal and Ara present in rhamnogalacturonan-I (RG-I) or Xyl part of xylogalacturonan (XG) were also detected in these fractions. High amounts of Xyl, especially in *C. tomentosa* and substantial amounts of Fuc and Man in all fractions indicate that further polysaccharides besides pectins are present in these fractions. A very unusual feature present mainly in the AmOx fractions is 3-*O*-methylgalactose, especially in *C. aspera*.

**FIGURE 1.**
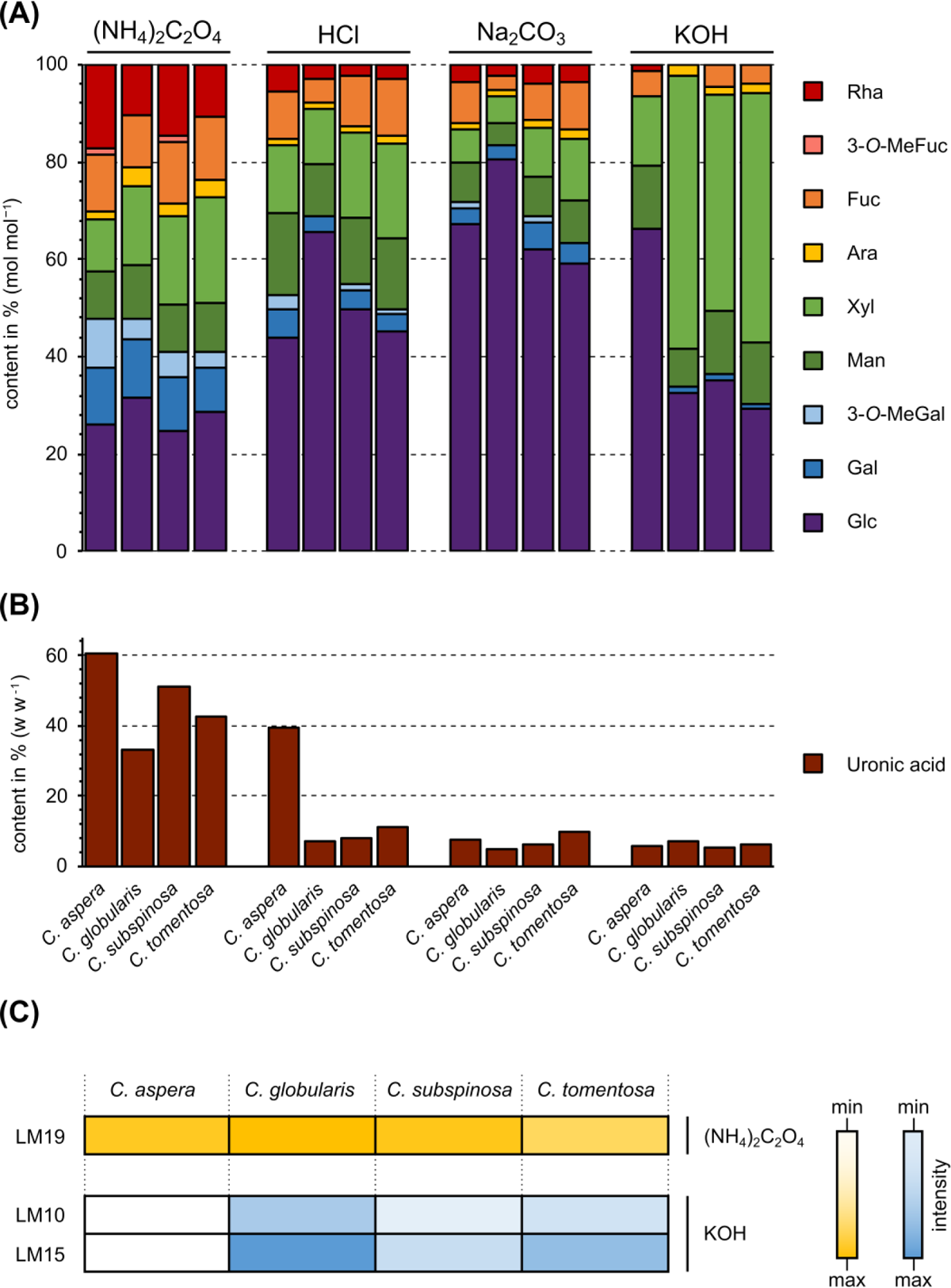
Monosaccharide composition of different cell wall fractions ((NH_4_)_2_C_2_O_4_, HCl, Na_2_CO_3_, KOH) from the streptophyte algae *C. aspera*, *C. globularis, C. subspinosa* and *C. tomentosa.* (a) Relative neutral monosaccharide composition determined by gas chromatography (GC; % mol mol^-1^). (b) Absolute content of uronic acids determined by colorimetric assay (w w^-1^ of dry fraction weight). (c) Reactivity of *Chara* sp. fractions with antibodies *via* ELISA after 40 min in the concentration 12.5 µg ml^-1^ ((NH_4_)_2_C_2_O_4_: LM19; KOH: LM10, LM15).

**TABLE 1.** Colorimetric determination of the content of uronic acids in the different cell wall fractions from different *Chara* species in % (w w^-1^).

|  | *C. aspera* | *C. globularis* | *C. subspinoso* | *C. tomentosa* |
| --- | --- | --- | --- | --- |
| AE | 5.8 | 3.4 | 3.6 | 4.2 |
| (NH<sub>4</sub>)<sub>2</sub>C<sub>2</sub>O<sub>4</sub> | 60.6 ± 1.7 | 33.1 ± 1.2 | 50.9 ± 2.5 | 42.8 ± 4.1 |
| HCl | 39.4 ± 8.5 | 7.1 | 8.2 | 11.3 |
| Na<sub>2</sub>CO<sub>3</sub> | 7.6 | 4.8 | 6.3 | 10.0 |
| KOH | 6.0 | 7.1 | 5.3 | 6.1 |

To confirm presence of pectins, the antibody LM19 raised against unesterified HG epitopes (Verhertbruggen *et al*., 2009b) was tested for binding affinity to the AmOx fractions (Figure 1) which revealed comparable amounts of unesterified HG in all *Chara* species. The antibody INRA-RU2 which recognizes the RG-I backbone showed no affinity to the AmOx fractions of all *Chara* species.

### Hemicellulose fractions of *Chara* sp. cell walls

In the alkaline fractions isolated with sodium carbonate (Table 2c), Glc was strongly dominating with 59% to 80%, while all other monosaccharides were present in amounts less than 10% (exception 12.7% Xyl in *C*. *tomentosa*), indicating occurrence of glucans.

**TABLE 2a.** Neutral monosaccharide composition of the ammonium oxalate fraction ((NH_4_)_2_C_2_O_4_) from different *Chara* species in % (mol mol^-1^ tr: trace value < 1%).

| Neutral monosaccharide | *C. aspera*<br>(NH<sub>4</sub>)<sub>2</sub>C<sub>2</sub>O<sub>4</sub><br>n=3 | *C. globularis*<br>(NH<sub>4</sub>)<sub>2</sub>C<sub>2</sub>O<sub>4</sub><br>n=3 | *C. subspinoso*<br>(NH<sub>4</sub>)<sub>2</sub>C<sub>2</sub>O<sub>4</sub><br>n=3 | *C. tomentosa*<br>(NH<sub>4</sub>)<sub>2</sub>C<sub>2</sub>O<sub>4</sub><br>n=3 |
| --- | --- | --- | --- | --- |
| 3-*O*-MeRha | tr | tr | tr | tr |
| Rha | 17.2 ± 0.3 | 10.4 ± 0.1 | 14.7 ± 0.2 | 10.8 ± 0.5 |
| 3-*O*-MeFuc | 1.2 ± 0.1 | tr | 1.1 ± 0.0 | tr |
| Fuc | 11.9 ± 0.1 | 10.8 ± 0.1 | 12.7 ± 0.1 | 12.9 ± 0.1 |
| Ara | 1.4 ± 0.0 | 3.9 ± 0.2 | 2.7 ± 0.1 | 3.5 ± 0.0 |
| Xyl | 10.7 ± 0.3 | 16.2 ± 0.2 | 18.0 ± 0.3 | 21.8 ± 0.2 |
| Man | 9.9 ± 0.4 | 10.8 ± 0.1 | 9.7 ± 0.1 | 9.9 ± 0.2 |
| 3-*O*-MeGal | 9.9 ± 0.3 | 4.2 ± 0.2 | 5.4 ± 0.1 | 3.5 ± 0.1 |
| Gal | 11.6 ± 0.1 | 12.1 ± 0.2 | 11.0 ± 0.2 | 8.9 ± 0.2 |
| Glc | 26.2 ± 0.7 | 31.6 ± 0.7 | 24.7 ± 0.3 | 28.7 ± 0.9 |

**TABLE 2b.**
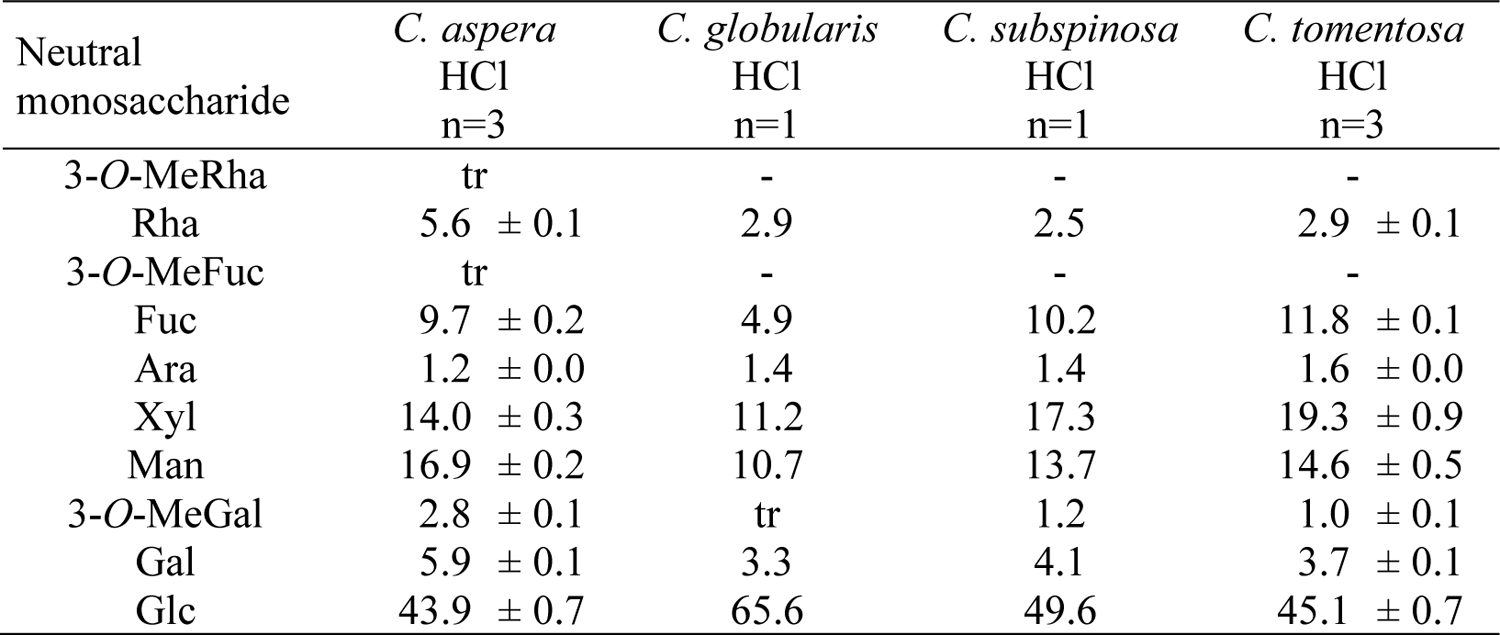
Neutral monosaccharide composition of the hydrochloric acid fraction (HCl) from different *Chara* species in % (mol mol^-1^_;_ tr: trace value < 1%).

| Neutral monosaccharide | *C. aspera*<br>HCl<br>n=3 | *C. globularis*<br>HCl<br>n=1 | *C. subspinoso*<br>HCl<br>n=1 | *C. tomentosa*<br>HCl<br>n=3 |
| --- | --- | --- | --- | --- |
| 3-*O*-MeRha | tr | - | - | - |
| Rha | 5.6 ± 0.1 | 2.9 | 2.5 | 2.9 ± 0.1 |
| 3-*O*-MeFuc | tr | - | - | - |
| Fuc | 9.7 ± 0.2 | 4.9 | 10.2 | 11.8 ± 0.1 |
| Ara | 1.2 ± 0.0 | 1.4 | 1.4 | 1.6 ± 0.0 |
| Xyl | 14.0 ± 0.3 | 11.2 | 17.3 | 19.3 ± 0.9 |
| Man | 16.9 ± 0.2 | 10.7 | 13.7 | 14.6 ± 0.5 |
| 3-*O*-MeGal | 2.8 ± 0.1 | tr | 1.2 | 1.0 ± 0.1 |
| Gal | 5.9 ± 0.1 | 3.3 | 4.1 | 3.7 ± 0.1 |
| Glc | 43.9 ± 0.7 | 65.6 | 49.6 | 45.1 ± 0.7 |

**TABLE 2c.**
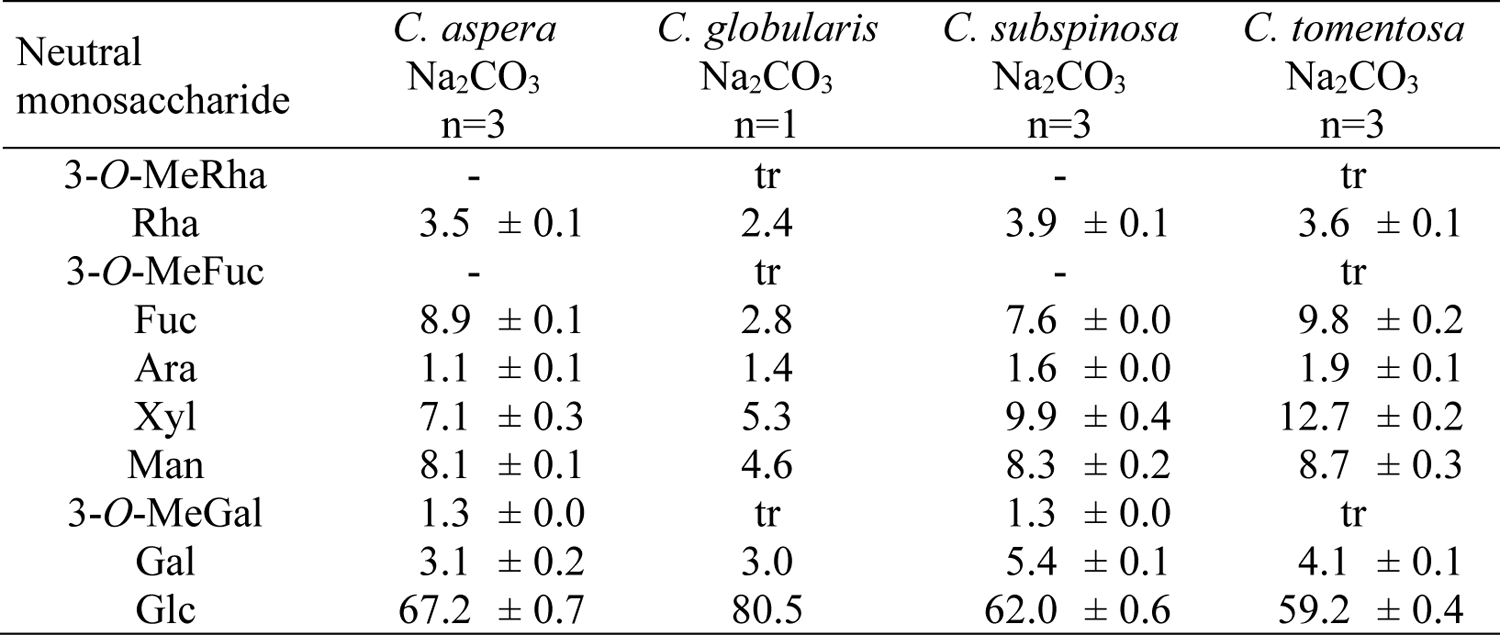
Neutral monosaccharide composition of the sodium carbonate fraction (Na_2_CO_3_) from different *Chara* species in % (mol mol^-1^ tr: trace value < 1%).

| Neutral monosaccharide | *C. aspera*<br>Na<sub>2</sub>CO<sub>3</sub><br>n=3 | *C. globularis*<br>Na<sub>2</sub>CO<sub>3</sub><br>n=1 | *C. subspinoso*<br>Na<sub>2</sub>CO<sub>3</sub><br>n=3 | *C. tomentosa*<br>Na<sub>2</sub>CO<sub>3</sub><br>n=3 |
| --- | --- | --- | --- | --- |
| 3-*O*-MeRha | - | tr | - | tr |
| Rha | 3.5 ± 0.1 | 2.4 | 3.9 ± 0.1 | 3.6 ± 0.1 |
| 3-*O*-MeFuc | - | tr | - | tr |
| Fuc | 8.9 ± 0.1 | 2.8 | 7.6 ± 0.0 | 9.8 ± 0.2 |
| Ara | 1.1 ± 0.1 | 1.4 | 1.6 ± 0.0 | 1.9 ± 0.1 |
| Xyl | 7.1 ± 0.3 | 5.3 | 9.9 ± 0.4 | 12.7 ± 0.2 |
| Man | 8.1 ± 0.1 | 4.6 | 8.3 ± 0.2 | 8.7 ± 0.3 |
| 3-*O*-MeGal | 1.3 ± 0.0 | tr | 1.3 ± 0.0 | tr |
| Gal | 3.1 ± 0.2 | 3.0 | 5.4 ± 0.1 | 4.1 ± 0.1 |
| Glc | 67.2 ± 0.7 | 80.5 | 62.0 ± 0.6 | 59.2 ± 0.4 |

**TABLE 2d.**
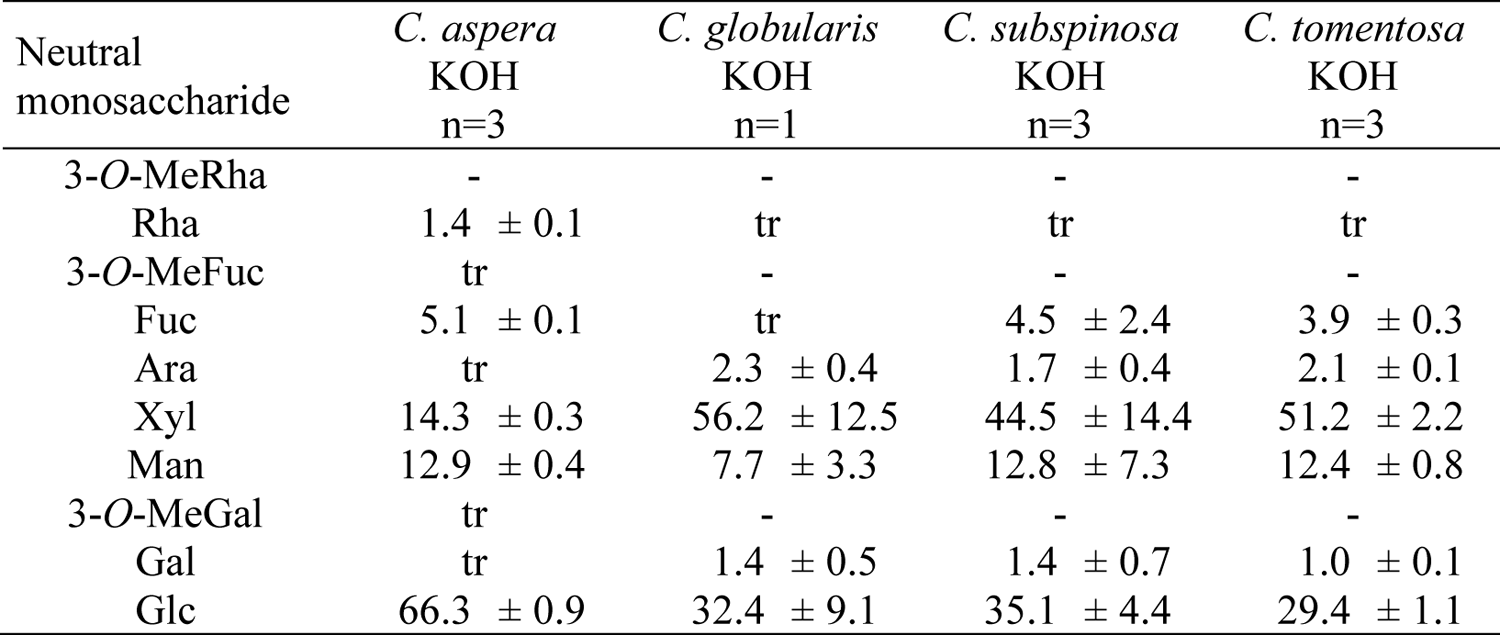
Neutral monosaccharide composition of the potassium hydroxide fraction (KOH) from different *Chara* species in % (mol mol^-1^ tr: trace value < 1%).

The KOH fractions (Table 2d) consisted mainly of Xyl and Glc, accompanied by Man. In all species, Xyl and Glc accounted for minimal 80% of the neutral monosaccharides, but the ratio of Xyl: Glc varied. Again, *C. aspera* was outstanding with a very low Xyl content in this fraction, and this was also confirmed by ELISA with the antibodies LM10 and LM15. These antibodies detect the epitopes of the non-reducing end of (1→4)-β-D-xylan (LM10; McCartney *et al*., 2005; Ruprecht *et al*., 2017) and XXXG-motif of xyloglucan (LM15; Marcus *et al*., 2008; Pedersen *et al*., 2012). *C. globularis* showed the highest absorption with both antibodies, followed by *C. tomentosa* and *C. subspinosa*. Interestingly, *C. aspera* revealed no binding to either antibody.

### Search for AGPs in *Chara* spp. cell walls

Water-soluble polysaccharides were precipitated with ethanol 80% (AE) and consisted mainly of Gal including 3-*O*-MeGal and Glc (Table S3a). The amounts of uronic acids in AE were quantified photometrically and varied between 3.4 % and 5.8 % (Table 1). In preliminary experiments, precipitation of AE with Yariv’s reagent yielded no AGP but resulted in a monosaccharide composition very similar to AE with slight increase in Gal content (data not shown), Therefore we used enzymatic treatments to purify a putative AGP which might be present in AE. As starch and some pectins might have been coextracted with these tentative AGPs, we treated AE fractions with α-amylase and pectinase, precipitated the remaining polysaccharides with ethanol 80% and dialyzed the precipitates to remove residual mono- or oligosaccharides (Table S3b). These treatments led to a decrease of Glc and an increase in Gal content in all species (compare Tables S3a and S3b. Comparable to the AmOx fraction, the highest amount of 3-*O*-MeGal with 19.3% was found in *C. aspera*. The monosaccharides Rha, Fuc, Xyl and Man were distributed in quite similar quantities between 7.5–16.5%, in all investigated *Chara* species, whereas the amounts of Ara were quite low (2.2-4.0%). Compared to the different *Chara* species, *Nitellopsis obtusa* was richer in Rha and Ara. To detect uronic acids, samples were reduced by use of sodium borodeuteride. There were only slight changes with increase of Gal and Glc (compare Tables S3b and 3). The mass spectra showed deuterated fragments deriving from GlcA and to a lesser extent, from GalA.

### Gel diffusion assay

A typical feature of AGPs is the precipitation with Yariv phenyl glycosides, like β-D-glucosyl Yariv-reagent (βGlcY). A gel diffusion assay provides a first hint whether AGPs are present in the sample. If AGPs are included, a red precipitation line will appear between the sample and the βGlcY (Clarke *et al*., 1979). The water-soluble fractions (AE) of all *Chara* species showed no Yariv precipitation, but after enzymatic treatment, weak precipitation lines were observable for all *Chara* species, but not for *Nitellopsis obtusa*, another member of the Charophyceae (Figure S1).

### Structure elucidation

The purified aqueous extracts (AE_AP) of all *Chara* species and also of *Nitellopsis obtusa* were subjected to linkage analysis (Table 4) to gain insight into polysaccharides present. Gal*p* as main monosaccharide was present as terminal residue (1.3-4.9 %), in 1,3-linkage (7.0-13.9 %), 1,6-linkage (5.7-43.5 %) and 1,3,6-linkage (5.0-13.5 %). Interestingly, these are the typical Gal linkage types present in AGPs. To identify linkage types of 3-*O*-MeGal, methylation was carried out by use of deuterated iodomethane. Mass spectra clearly revealed that some of the terminal and also 1,6-linked Gal*p* residues are present as 3-*O*-MeGal in the native samples. Typical AGP linkage types of Ara*f* (terminal and 1,5-linked Ara) worth mentioning were present only in *Nitellopsis* AE_AP. All samples and especially that of *Nitellopsis* were further characterized by high amounts of 1,4-linked Rha*p,* accompanied by small amounts of other linkage types of this monosaccharide. The heterogeneity of the sample with many different monosaccharides was confirmed with Glc/GlcA, Fuc and Xyl as terminal residues and further linkage types of desoxyhexoses (Rha and/or Fuc) and hexoses (Man and Glc). The mass spectrum of terminal Glc in the uronic acid reduced sample clearly revealed presence of GlcA in the native sample due to deuterated fragments originating from sodium borodeuteride used for reduction. The peak of 1,4-Hexp also contained small amounts of deuterated fragments, thus revealing the presence of 1,4-GalA or 1,4-GlcA (same retention time) in the native sample.

### ELISA with antibodies directed against AG glycan epitopes

To detect arabinogalactan-protein glycan epitopes, the four AE fractions were tested for binding affinities to the antibodies KM1, JIM13, LM2 and LM6 (Figure 2).

**FIGURE 2.**
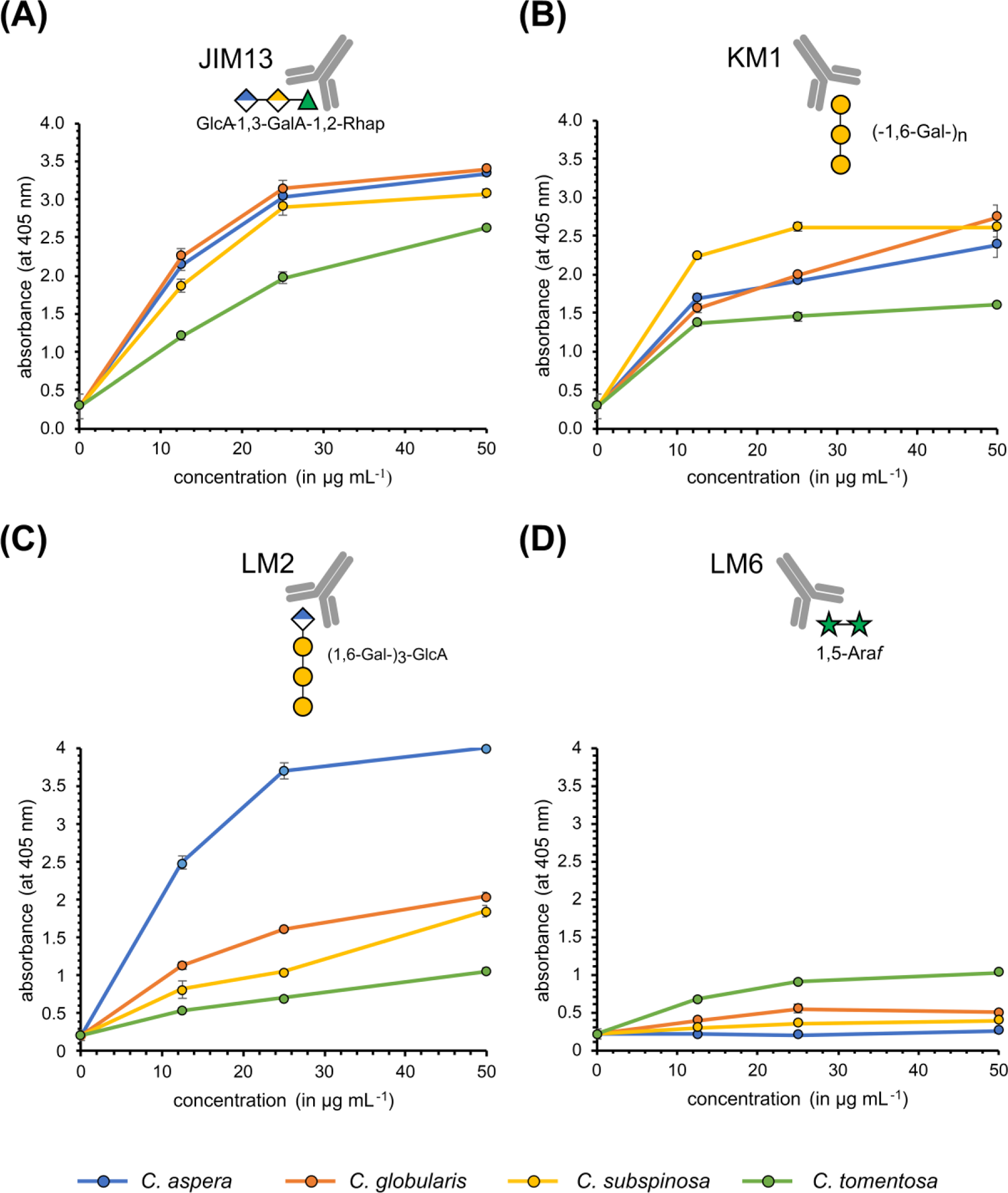
Reactivity of monoclonal antibodies with polysaccharide structures in AEs of different *Chara* species determined *via* ELISA. Antibodies are directed against typical AGP glycan structures, which are briefly symbolized above each diagram. (A) JIM13; (B) KM1; (C) LM2; (D) LM6. For epitopes of the antibodies, see Table S1.

The antibody KM1 (Figure 2B) detects the (1→6)-β-D-Gal*p* units in AGs type II (Classen *et al*., 2004; Ruprecht *et al*., 2017). For JIM13 (Figure 2A), the exact epitope is still under debate, but it seems obvious that this antibody recognizes AGPs and that Rha and uronic acids are part of the epitope (Pfeifer *et al*., 2022). The trisaccharide β-D-GlcA*p*-(1→3)-α-D-GalA*p*-(1→2)-α-L-Rha has been shown to bind to JIM13 (Yates *et al*., 1996). The affinity of these antibodies to AE of all *Chara* species were strong, concentration dependent and comparable between the species with slightly weaker binding of the AE of *C. tomentosa*.

The antibody LM2 (Smallwood *et al*., 1996, Figure 2C) recognizes (1→6)-β-D-Gal*p* units with terminal ß-D-GlcA*p* present in AGPs (Ruprecht *et al*., 2017) and showed moderate binding properties to AE of *C. globularis*, *C. subspinosa* and *C. tomentosa*, but very strong reactivity with AE of *C. aspera*, which contains highest amounts of 1,6-linked Gal (Table 4).

The antibody LM6 (Figure 2D) identifies (1→5)-α-L-Ara*f* oligomers in arabinans or AGPs (Verhertbruggen *et al*., 2009a). The AE of *C. tomentosa* showed very weak binding to LM6, whereas the other *Chara* species showed no reactivity. Low or no affinity to this antibody is in accordance with low Ara content in AE of these species (Table 4).

### Quantification of hydroxyproline

In AGPs, several arabinogalactan moieties are covalently linked to a protein backbone *via* the amino acid hydroxyproline. According to Stegemann and Stalder (1967), the hydroxyproline contents of all *Chara* AE samples were quantified photometrically. The results were very interesting, as hydroxyproline was not detectable in any of the samples. Samples (AE) of *C. globularis*, *C. subspinosa* and *C. tomentosa* collected in Austria as well as from *N. obtusa* were also free of Hyp. For this reason, we were interested whether prolyl-4-hydroxylases are present in the genome of *Chara braunii*.

### Bioinformatic search for Prolyl-4-hydroxylase

BLAST search or the available genome data of *Chara braunii* revealed presence of seven putative P4H homologs. In a detailed analysis of the aligned sequences, absence or presence of relevant amino acids in the oxoglutarate-binding site as well as the iron-binding region (according to Hieta & Myllyharju, 2002; Keskiaho *et al*., 2007; Koski *et al*., 2007; Koski *et al*., 2009) was investigated (Table S4a and S4b). Only in two cases all important residues were found (GBG_70085 and GBG_85197). One of these protein sequences (GBG_70085) lacks Arg^93^ (nomenclature based on *C. reinhardtii* P4H), which most likely leads to complete inactivation of P4H activity (Koski *et al*., 2007). Therefore, only one sequence (GBG_85197) was used for protein structure modelling *via* SWISS-MODEL (Figure 3).

**FIGURE 3.**
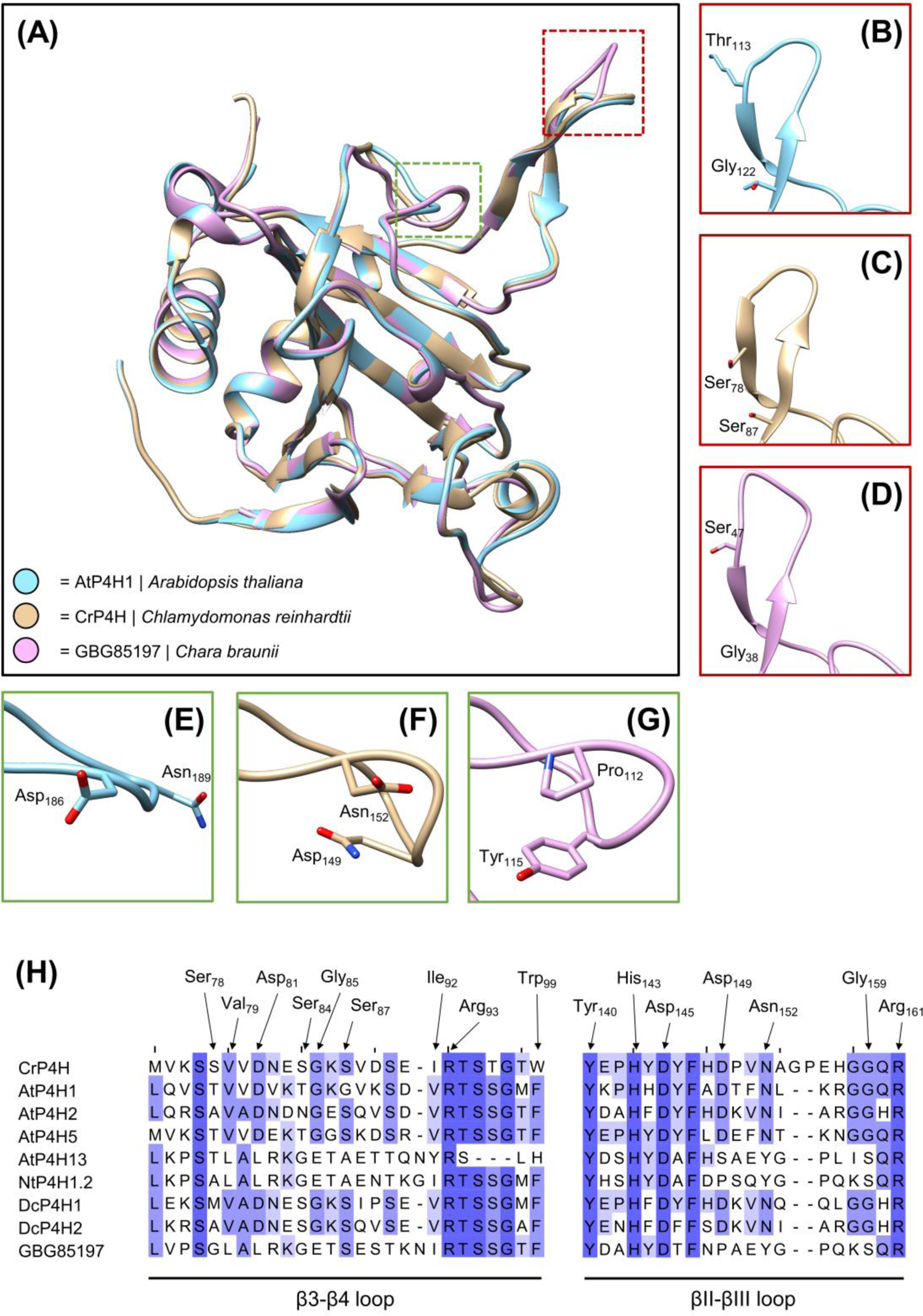
Modelled protein structures of *Chara braunii* prolyl-4-hydroxylase (P4H) and two functionally described P4Hs of *Arabidopsis thaliana* and *Chlamydomonas reinhardtii*. (A) Overlay of the three (modelled) structures with highlighted regions within the substrate binding tunnel (red: β3-β4 loop; green: βII-βIII loop). (B-D) Detailed comparison of the β3-β4 loop. (E-G) Detailed comparison of the βII-βIII loop. (H) Alignment of *C. braunii* P4H with functionally described P4Hs. Only the regions shown in panels A-G are presented and relevant amino acids are highlighted with boxes.

The overlaid protein structures (Figure 3A) revealed overall matching topology in nearly all parts of the molecules but showed diverging structures in the putative substrate binding channel of *C. braunii* P4H in comparison to AtP4H1 and CrP4H (see red and green boxes in Figure 3H). Some of these structural differences (Pro instead of Asp^149^; Tyr instead of Asn^152^) are also visible in some of the P4H sequences used for comparison (AtP4H13 and NtP4H1.2). Both substrate channel regions, the β3-β4 loop as well as the βII-βIII loop, contain flexible parts which show a low degree of alignment (Figure 3B-H) flanked with phylogenetically highly conserved amino acids.

## DISCUSSION

### Polysaccharides of Chara cell walls

Although taxonomically defined differences in cell wall composition become more and more obvious, knowledge on streptophyte algal cell walls is still limited. Although there is some evidence for the occurrence of nearly all seed plant polysaccharides in streptophyte algae (Domozych & Bagdan, 2022), recent studies are often based on immunocytochemistry which has limitations as epitopes of antibodies are not always clearly defined. Thus, isolation and analytical characterization of cell wall polysaccharide fractions of streptophyte algae as closest living relatives to land plants helps to understand cell wall evolution during plant terrestrialization.

### Pectin composition in Charales hints towards presence of (substituted) HGs but absence of RG-I

For isolation of pectic fractions, we used standard procedures with chelators and hot acid (Scheller & Ulvskov, 2010). The high amount of the AmOx fraction in *C. aspera* was striking (21.2% of dry plant material, compared to 2.3 – 5.9% in the other *Chara* species). Since pectins are characterized by a high content of GalA, the high amounts of uronic acids in the two pectic fractions in all *Chara* species were not surprising, but again extraordinary in *C. aspera* with 60% in the AmOx and 40% in the HCl fraction. As *C. aspera* is the only investigated species which is salt tolerant, the involvement of pectins in adaptation to salt water is likely. High concentrations of uronic acids were also detected in four other members of the Charophyceae, *Chara corallina*, *Chara vulgaris, Nitella flexilis* and *Nitellopsis obtusa* (O’Rourke *et al*., 2015; Popper & Fry, 2003; Pfeifer *et al*., 2022) as well as in *Coleochaete scutata* (Coleochaetophyceae). In contrast, the pectic fractions of *Spirogyra pratensis* (Zygnematophyceae) contained only low amounts of uronic acids (Pfeifer *et al*., 2022) and those of two *Klebsormidium* species (Klebsormidiaceae) were completely free of GalA and also GlcA (O’Rourke *et al*., 2015), thus revealing strong differences in composition of pectic fractions of streptophyte cell walls.

Pectins mainly consisting of non-esterified GalA have been purified from *Nitella translucens* and *Chara australis* (Anderson & King, 1961a, 1961b). Immunocytochemistry with antibodies directed against HG (LM19, LM20, JIM5, JIM7, 2F4) confirmed presence of this pectic polysaccharide in many members of streptophyte algae (Proseus & Boyer, 2006; Eder & Lütz-Meindl, 2008, 2010; Domozych *et al*., 2014; Herburger *et al*., 2019; Palacio-López *et al*., 2019; Permann *et al*., 2021a, 2021b; Sørensen *et al*., 2011), but glycan microarray with HG antibodies JIM5 and 2F4 also revealed strong differences. Whereas there was strong reactivity with CDTA extracts of *Chara*, *Coleochaete*, *Cosmarium*, *Penium* and *Netrium*, **no** reactivity was detected in *Spirogyra*, *Klebsormidium* and *Chlorokybus* (Sørensen *et al*., 2011). In contradiction to these results, CDTA extracts of *Spirogyra mirabilis* and *Mougeotia disjuncta* showed strong binding especially to LM19 and JIM5, which both recognize HG with low degree of esterification (Permann *et al*., 2021a, 2021b). To conclude, it seems that presence of homogalacturonans is an ancient feature of streptophytes, although secondarily lost in some species.

In case of presence of RG-I, Rha, Gal and Ara should be present in the pectic fractions. Rha accounts for over 10% of the neutral monosaccharides in the AmOx fractions of all *Chara* species investigated in this study and Gal (including3-*O*-MeGal) is also present in higher amounts, but the content of Ara is low. Low amounts of Ara were also verified for AmOx fractions of *C. vulgaris* and *N. flexilis* (O’Rourke *et al*., 2015). In *Chara australis* and several members of the Zygnematophyceae, linkage types typically associated with Ara and Gal sidechains of RG-I were absent or present at low levels (Sørensen *et al*., 2011). Isolation of pectins by oxalate followed by paper chromatography suggested presence of RG I only in the later-diverging CGA *Nitella flexilis*, *Chara vulgaris* and *Coleochaete scutata* (O’Rourke *et al*., 2015). The antibody LM5 directed against an epitope containing 1,4-linked Gal present in RG-I showed weak binding to CDTA extracts from *Mougeotia disjuncta* and *Spirogyra mirabilis* (Permann *et al*., 2021a, 2021b) but other authors detected no binding of this antibody to CDTA extracts from *Chara corallina*, *Coleochaete nitellarum* and several members of the Zygnematophyceae (Sørensen *et al*., 2011). To conclude, presence of RG-I in streptophyte algae is still not finally settled but more likely for members of the ZCC grade. Presence in two members of the Zygnematophyceae has been confirmed by binding of the antibodies INRA-RU1 and 2, which recognize the RG-I backbone, to CDTA extracts from *Mougeotia* and *Spirogyra* (Permann *et al*., 2021a, 2021b). In contrast, the AmOx fractions of all investigated *Chara* species showed no binding to INRA-RU2, suggesting that RG-I is not present at least in this genus of the Charophyceae.

A special feature of all pectic fractions of the four *Chara* species was 3-*O*-MeGal, which was also detected in *Chara vulgaris* and *Chara corallina* (Sørensen *et al*., 2011). It has been suggested that this methylated monosaccharide might have replaced Ara in RG-I of *Chara* species (O’Rourke *et al*. 2015), which is contradictory to our results which propose absence of RG-I in *Chara* species.

Investigations on glycosyl linkages revealed none of the characteristic sugars of RG II (2-*O*MeFuc, 2-*O*MeXyl, apiose, aceric acid) in streptophyte algae (Sørensen *et al*., 2011), thus further investigations are also necessary to clarify the evolutionary roots of RG II.

### Ancestral hemicellulose core structures seem to have diversified after the divergence of Charales and all other streptophytes

Since most hemicelluloses can be extracted under alkaline conditions (Scheller & Ulvskov, 2010), two alkaline fractions (sodium carbonate and KOH) were extracted from the cell walls of the four investigated *Chara* species. Xyl, Man, and Glc were abundant in the hemicellulose fractions of all *Chara* sp., especially Xyl and Glc in the KOH fractions, which might be part of xylans, xyloglucans and (gluco)mannans. Striking was the low Xyl content of the KOH fraction of *C. aspera*. This is also demonstrated by the Xyl:Glc ratio. In the other *Chara* species, this ranges from 1.3 – 1.7:1, whereas *C. aspera* shows a ratio of 1:4.6. Presence of xylans and xyloglucans in the cell walls of *C. globularis*, *C. subspinosa* and *C. tomentosa* was confirmed by ELISA with LM10 (xylans) and LM15 (xyloglucans). As the KOH fraction of *C. aspera* showed no reactivity with both antibodies, this species might be characterized by lack or structural modifications of both hemicelluloses. It has to be mentioned, that LM10 only detects non or low substituted xylans (McCartney *et al*., 2005). Interestingly, the NaOH extract of *Chara corallina* also showed no binding to LM10, but bound to LM11, which detects arabinoxylans as well (Sørensen *et al*., 2011).

In general, today there is good evidence for presence of xylans and xyloglucans in streptophyte algae. This is based on investigations with antibodies directed against xylan and xyloglucan epitopes (Eder & Lütz-Meindl, 2008; Domozych *et al*., 2009; Sørensen *et al*., 2011; Ikegaya *et al*., 2012; Mikkelsen *et al*., 2021; Permann *et al*., 2021a, 2021b) but also on structural characterization (Ikegaya *et al*., 2008; Sørensen *et al*., 2011; Mikkelsen *et al*., 2021) and investigations on enzymes involved in biosynthesis (Del-Bem & Vincentz, 2010; Franková & Fry, 2021; Mikkelsen *et al*., 2014; Mikkelsen *et al*., 2021, Nishiyama *et al*., 2018).

Presence of fucosylated and galactosylated xyloglucans has been shown for *Mesotaenium* (Mikkelsen *et al*., 2021), underlining that these xyloglucan substitutions known from land plants are also present at least in some streptophyte algae. Especially in the sodium carbonate fractions of the four *Chara* species investigated here, Gal and Fuc are present in amounts between 2.8% and 9.8%, maybe indicating fucosylated and/or galactosylated xyloglucans.

### Purified aqueous extracts include unusual rhamnans and galactans

#### Gel diffusion assay

Detection of AGPs is possible by precipitation with βGlcY (Yariv *et al*., 1962). Although the exact precipitation mechanism has not yet been precisely determined, it is certain that a β-1,3-linked galactan is important for this interaction (Kitazawa *et al*., 2013; Paulsen *et al*., 2014). In addition, it has been proposed that high concentrations of 1,6-linked galactans can interfere with the precipitation, either directly (Kiyohara *et al*., 1989) or relatively (Kitazawa *et al*., 2013; Paulsen *et al*., 2014).

In a radial gel diffusion assay with βGlcY, AE of all *Chara* species showed no precipitation lines, proposing absence of AGPs in this genus. After purification of AE by treatment with α-amylase and pectinase (AE_AP; deletion of starch and homogalacturonan, which might have been coextracted) small precipitation lines could be detected (Figure S1), possibly originating from 1,3-linked galactans and not from AGPs, as no Hyp was present (see below).

#### Analyses of the carbohydrate moiety

Seed plant, fern and bryophyte AGPs have been isolated from aqueous extracts (AE) rich in Ara and Gal by precipitation with βGlcY (Bartels & Classen, 2017; Classen *et al*., 2004; Happ & Classen, 2019; Mueller *et al*., 2023). The AE fraction compositions of the cell walls of the investigated *Chara* species were different, with high Gal contents Gal but only low amounts of Ara. AE of *Nitellopsis* was slightly different with higher amounts of Rha and Ara. AE of *C. aspera* was outstanding due to the high content of uronic acids (over 30%). Furthermore, the presence of 3-*O*-MeGal in all AE, especially in *C. aspera*, was extraordinary. Methylation of one hydroxyl-group of monosaccharides is a rare modification of different carbohydrates and has been described for some bacteria, fungi, worms, molluscs and also plants (for review see Staudacher, 2012). It offers the possibility to modulate glycan structures, thereby leading to new biological activities. In plants, galactose is the most common methylated hexose (Staudacher, 2012), especially in algae. 4-*O*-MeGal and 3-*O*-MeGal occur in red algae (Araki *et al*., 1986; Allsobrook *et al*., 1974; Navarro & Stortz, 2008). In cell walls of chlorophytic green algae, polysaccharides containing 3-*O*-MeGal, 2-*O*-MeRha and 3-*O*-MeRha (*Chlorella vulgaris*) or 3-*O*-MeRha and 3-*O*-MeFuc (*Botryoccoccus braunii*) were found (Allard & Casadevall, 1990; Ogawa *et al*., 1994; Ogawa *et al*., 1997). For *Chlorella vulgaris*, it was shown that 3-*O*-MeGal is part of a neutral galactan with a ratio of Gal: 3-*O*-MeGal of 7.1:1 (Ogawa *et al*., 2001). In *Spirogyra pratensis*, a Zygnematophyceae, 3-*O*-MeGal, 2-*O*-MeRha and 3-*O*-MeRha were detected only in traces (Pfeifer *et al*., 2022). In the streptophyte algae *Klebsormidium flaccidum, Chara corallina* and *Coleochaete scutata* 3-*O*-MeRha was also identified in small amounts (Popper & Fry 2003), but 3-*O*-MeGal was not detected at all (Popper *et al*., 2001). More recently it was shown that pectins of *Chara vulgaris* are characterized by the presence of 3-*O*-MeGal (ÓRourke *et al*., 2015). Interestingly, among the land plants, the cell walls of bryophytes and ferns contain 3-*O*-MeRha, especially as part of AGPs (Bartels *et al*., 2017; Bartels & Classen, 2017 Happ & Classen, 2019, Baumann *et al*., 2021, Mueller *et al*., 2023), or RG-II (Matsunaga *et al*, 2004?), whereas higher concentrations of 3-*O*-MeGal were detected in the cell walls of the young leaves of both homosporous (*Lycopodium*, *Huperzia* and *Diphasiastrum*) and heterosporous (*Selaginella*) lycophytes (Popper *et al*., 2001). Although only rarely, 3-*O*-MeGal was found in some seed plant polysaccharides as well (Barsett & Paulsen, 1992; Hokputsa *et al*., 2004), even as part of an arabinogalactan-protein (Capek, 2008).

To verify the presence of arabinogalactans, antibodies directed against arabinogalactan epitopes present in AGPs are often used, e.g. JIM8, JIM13, JIM16, LM2, LM6, LM14, KM1, MAC207. It should be taken into account that the exact epitopes of some of these antibodies are still unknown. We used JIM13, LM2, LM6 and KM1, and AE of all *Chara* species investigated showed comparable binding profiles in ELISA. There was nearly no binding of Chara AE to LM6, indicating lack of 1,5-linked Ara. In contrast, AE of *Nitellopsis obtusa* bound to LM6 and structure elucidation revealed high amounts of 1,5-linked Ara (Table 4). KM1, which recognizes 1,6-linked Gal, bound to AE of all *Chara* species and also to AE of *Nitellopsis* (Pfeifer *et al*., 2022) thus revealing presence of 1,6-linked Gal. Binding of AE to JIM13 with an epitope probably including Rha and uronic acids was comparable for the *Chara* species but very low for *Nitellopsis* (Pfeifer *et al*., 2022). In ELISA with LM2, *C. aspera* was outstanding. According to Ruprecht *et al*. (2017), this antibody recognizes (1→6)-β-D-Gal*p* units with terminal ß-D-GlcA*p*. Probably the high content of 1,6-Gal residues in AE of *C. aspera* is responsible for the higher affinity to this antibody. In ELISA experiments with AE of *Spirogyra pratensis* (Pfeifer *et al*., 2022), no affinity was detected for LM6 and KM1, whereas there was strong binding to JIM13. CDTA extracts of other members of the Zygnematophyceae (*Spirogyra mirabilis* and *Mougeotia sp*.) also showed strong interaction with JIM13 (Permann *et al*., 2021a, 2021b). In glycan microarray, different antibodies directed against AG epitopes (LM2, LM14, JIM8, JIM13 and MAC207) have been tested for interaction with CDTA extracts of different streptophyte algae (Sørensen *et al*., 2011). All antibodies (except for LM2) and especially JIM13 bound to extracts of *Chara corallina*, but JIM13 was the only antibody binding to *Spirogyra* extracts, thus confirming strong structural differences between the Charophyceae and the Zygnematophyceae. Microscopic investigations also revealed positive reaction of JIM13 with several members of the Zygnematophyceae (Eder *et al*., 2008; Domozych *et al*., 2009; Ruiz-May *et al*., 2018; Palacio-Lopez *et al*., 2019). Interestingly, *Klebsormidium flaccidum*, representing the lower-branching KCM-grade, showed no reactivities with antibodies directed against arabinogalactan epitopes (Sørensen *et al*., 2011, Steiner *et al*., 2020).

Structural analyses verified presence of unusual galactans. Linkage types typical for AGPs were present and also detected by antibodies directed against AGPs (see above). 1,3-linked Gal chains are possibly responsible for the weak interaction with Yariv’s reagent (Kitazawa *et al*., 2013; Paulsen *et al*., 2014). A special feature were high amounts of 3-*O*-MeGal, present in 1,6-linkage or as terminal residue. The ratio of Gal to 3-*O*-MeGal varied (compare Table 3) and was highest for *C. aspera* (1 : 1.2). In the other species, Gal was dominant (1 : 0.5 for *C. subspinosa*, 1 : 0.4 for *C. globularis*, 1:0.3 for *C. tomentosa* and 1 : 0.1 for *N. obtusa* AE_AP_UR. Furthermore, the relations of the different linkage types were different between the species, for structural proposals see Figure 4. *C. aspera* is outstanding within the Chara species due to longest 1,6-Gal side chains, whereas *N. obtusa* is special due to short 1,6-linked Gal side chains. A galactan with similar structure has been isolated from *Chlorella vulgaris*, a unicellular member of the Chlorophyta (Ogawa *et al*., 2001), indicating that this polysaccharide has been conserved in the Charales. The structure of this *Chlorella* galactan is most comparable to that of *C. aspera* with even longer 1,6-linked Gal side chains, but with a lower degree of methylated Gal (Ogawa *et al*., 2001).

**FIGURE 4.**
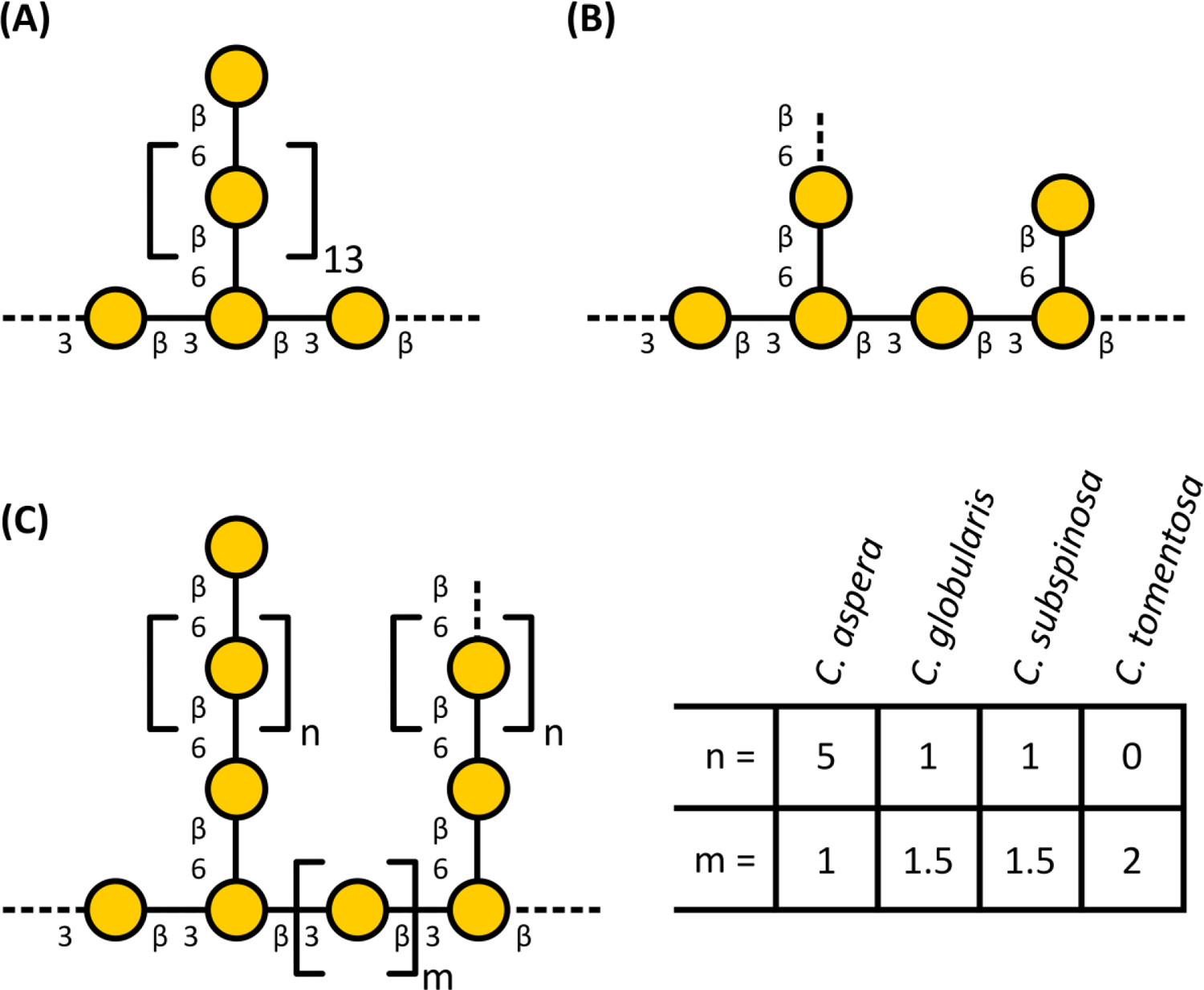
Partial structural proposals for galactans of *Chara* species in the context of other algal galactan structures. (A) Structural proposal of *Chlorella vulgaris* based on the published results of Ogawa *et al*. (2001). Patterns of methylation are not shown. (B) Galactan structures of *Nitellopsis obtusa*. (C) Galactan structures of the investigated four *Chara* species. The structural features were inferred from the linkage-type analysis. The dashed lines symbolize attachment sites for galactoses (main chain) or for other monosaccharides (side chain). Part of the Gal residues are methylated (see Table 3 for amounts of 3-*O*-MeGal).

**TABLE 3.** Neutral monosaccharide composition of dialyzed water-soluble polysaccharides, pre-treated with amylase and pectinase after reduction of uronic acids (AE_AP_UR), from different *Chara* species and *Nitellopsis obtusa* in % (mol mol^-1^_;_ tr: trace value < 1%).

| Neutral mono-saccharide | *C. aspera*<br>AE_AP_UR<br>n=3 | *C. globularis*<br>AE_AP_UR<br>n=3 | *C. subspinoso*<br>AE_AP_UR<br>n=3 | *C. tomentosa*<br>AE_AP_UR<br>n=3 | *N. obtusa*<br>AE<sub>purified</sub>_AP_UR<br>n=3 |
| --- | --- | --- | --- | --- | --- |
| Rha | 12.6 ± 0.1 | 10.7 ± 0.2 | 13.7 ± 0.2 | 10.0 ± 0.2 | 18.6 ± 0.3 |
| 3-*O*-MeFuc | - | - | 1.9 ± 0.1 | - | - |
| Fuc | 8.7 ± 0.0 | 14.1 ± 0.1 | 8.7 ± 0.2 | 11.6 ± 0.1 | 6.4 ± 0.0 |
| Ara | 2.1 ± 0.1 | 3.7 ± 0.1 | 2.3 ± 0.1 | 2.1 ± 0.1 | 14.4 ± 0.1 |
| Xyl | 6.7 ± 0.4 | 8.6 ± 0.1 | 9.7 ± 0.0 | 12.8 ± 0.8 | 5.1 ± 0.0 |
| Man | 9.0 ± 0.1 | 9.9 ± 0.1 | 10.7 ± 0.2 | 12.8 ± 0.2 | 8.0 ± 0.2 |
| 3-*O*-MeGal | 21.3 ± 0.3 | 8.1 ± 0.2 | 10.1 ± 0.1 | 4.7 ± 0.2 | 2.5 ± 0.0 |
| Gal | 18.3 ± 0.2 | 22.4 ± 0.2 | 20.2 ± 0.4 | 16.7 ± 0.1 | 26.8 ± 0.0 |
| Glc | 21.3 ± 0.7 | 22.5 ± 0.2 | 22.7 ± 0.2 | 29.3 ± 0.4 | 18.2 ± 0.1 |

**TABLE 4.** Linkage type analysis of AE_AP from *Chara* spp. and *N. obtusa* in % (w w^-1^, n=1).

| Mono-saccharide | linkage type | *C. aspera* | *C. globularis* | *C. subspinoso* | *C. tomentosa* | *N. obtusa* |
| --- | --- | --- | --- | --- | --- | --- |
|  |  | AE_AP | AE_AP | AE_AP | AE_AP | AE_AP |
| *Galp* | 1,3,6- | 7.0 | 5.7 | 6.0 | 4.5 | 13.2 |
|  | 1,3- | 6.7 | 9.0 | 9.7 | 9.0 | 13.6 |
| *Galp*/<br>*3-O-MeGalp* | 1,6- | 41.6 | 13.9 | 11.1 | 5.1 | 10.0 |
|  | 1- | 4.2 | 4.4 | 2.8 | 2.7 | 1.3 |
| *GlcP* | 1,3- | 1.0 | 4.1 | 3.7 | 5.9 | 2.1 |
|  | 1- | 5.1 | 7.7 | 8.4 | 11.8 | 2.6 |
| *Hexp* | 1,2,4,6- | 1.8 | 2.8 | 5.0 | 3.6 | - |
|  | 1,4,6- | - | 2.4 | 2.2 | 1.7 | 1.1 |
|  | 1,3,4- | - | 3.2 | 2.4 | 4.9 | 1.0 |
|  | 1,2,6- | - | 3.4 | 5.0 | 7.5 | 4.3 |
|  | 1,4- | 3.3 | 7.3 | 6.2 | 6.4 | 4.4 |
|  | 1,2- | 3.1 | 5.2 | 4.7 | 5.5 | 2.1 |
| *Rhap* | 1,2,4- | - | 1.0 | tr | tr | 1.0 |
|  | 1,4- | 15.5 | 9.1 | 12.4 | 10.7 | 19.7 |
|  | 1,2- | 1.4 | 1.8 | 1.4 | 1.5 | 0.0 |
|  | 1- | 1.9 | 2.3 | 2.6 | 2.2 | 4.0 |
| *Fucp* | 1- | 4.4 | 5.8 | 9.3 | 7.2 | 3.4 |
| *Desoxyhexp* | 1,3,4- | - | 3.1 | tr | tr | - |
|  | 1,3- | - | 1.7 | 1.3 | 1.8 | 1.3 |
| *Araf* | 1,5- | - | - | - | - | 5.6 |
|  | 1- | 1.2 | 1.6 | 1.3 | 1.2 | 6.8 |
| *Xylp* | 1- | 1.8 | 4.5 | 4.5 | 7.1 | 2.5 |

Another interesting finding of linkage-type analysis were high amounts of 1,4-linked Rha*p* in all investigated members of the Charales. To the best of our knowledge, no plant polysaccharide with a substantial amount of 1,4-linked Rha*p* has been described up to now. Typical linkage types known for Rha in seed plants are 1,2- and 1,2,4-linked Rha residues as part of RG-I. A main component of the fibers extracted from the green algae *Monostroma angicava* and *Monostroma nitidum* (Chlorophyceae) are sulfated rhamnans with 1,3- and 1,2-linked Rha residues (Liu *et al*., 2018; Suzuki & Terasawa, 2020) and the rhamnogalactan-protein isolated from *Spirogyra pratensis* is rich in 1,3-linked Rha*p* (Pfeifer *et al*., 2022).

#### Hydroxyproline seems to be absent in Charales and AGP-associated P4H activity is questionable

The protein backbone of classical AGPs consists of an *N*-terminal signal peptide, a protein sequence rich in proline (Pro), alanine (Ala), serine (Ser), and threonine (Thr; PAST), and a C-terminal domain (Johnson *et al*., 2018). The carbohydrate moieties are covalently linked to the protein part *via* Hyp, which is synthesized by the hydroxylation of Pro to Hyp, catalyzed by the enzyme prolyl 4-hydroxylase (P4H; Seifert *et al*., 2021).

In different studies, the occurrence of Hyp was tested in various chlorophyte algae and the charophyte algae *Spirogyra pratensis*, *Nitella* sp. and *Nitellopsis obtusa*. Hyp was detected in all algae, except of the two algae of the Charophyceae family: *Nitella* and *Nitellopsis* (Gotelli & Cleland, 1968; Morrison *et al*., 1993; Pfeifer *et al*., 2022; Thompson & Preston, 1967). Our results that Hyp is missing in all investigated *Chara* species is in support of these previous results and fosters the assumption that Hyp is absent in all members of the Charophyceae.

At the first glimpse, the finding of seven P4H homologs *via* BLAST analysis is in contrast to this statement. With the here presented analysis of the relevant amino acids in both catalytic sites (compare Hieta & Myllyharju, 2002; Keskiaho *et al*., 2007; Koski *et al*., 2007; Koski *et al*., 2009) it is likely that most homologs lost their functionality for prolyl-hydroxylation either in general or with regard to the characteristic HRGP substrates. The substrate specificity is highly dependent on the substrate tunnel being shaped by the β3-β4 loop as well as the βII-βIII loop, which cover the tripeptide structure of the substrate. In direct comparison with various vertebrate P4H substructures, it was shown that both regions have a certain degree of flexibility (Koski *et al*., 2009). Numerous studies with recombinant enzyme expression showed that substrate structures resemble this high flexibility (Mócsai *et al*., 2021; Velasquez *et al*., 2015; Vlad *et al*., 2010). Despite of the described activities for P4Hs on different substrates, it is not possible to predict specificity conclusively (compare Moussu & Ingram, 2023). The few studies comparing substrate activities (Mócsai *et al*., 2021; Velasquez *et al*., 2015; Vlad *et al*., 2010) have included an AGP substrate in their analyses but also reveal that for *C. reinhardtii* P4H this activity was not determined (Vlad *et al*., 2010). The selected candidate sequence GBG85197 from *C. braunii* showed overall similarity to the two P4Hs from *A. thaliana* and *C. reinhardtii* (Figure 3) with various structural variations in the two flexible regions. Even though the catalytically relevant amino acids are present (Hieta & Myllyharju, 2002; Keskiaho *et al*., 2007; Koski *et al*., 2007; Koski *et al*., 2009) this could mean extremely different or absent activity. Timmins *et al*. (2017) performed excessive mutations in the two investigated loops and showed inactivity, changed regioselectivity or different substrate activity by mutation of single amino acids. Presence of a putatively functional P4H and lack of hydroxyproline in the biochemical assays is explainable in different ways: (i) The occurrence of the coding gene in the genome is not equivalent to the translation to a functional protein which could be occasionally expressed or not expressed following a distinct (environmental) stimulus. (ii) The P4H coding gene could be a remnant of the least common ancestor of all Chloroplastida, which utilized the enzyme for hydroxylation of very specialized substrates. Exemplarily, the cell walls in Chlamydomonales are formed nearly exclusively by (glyco-)proteins showing *O*-glycosylation (Bollig *et al*., 2007; Ferris *et al*., 2001; Miller *et al*., 1972; Voigt *et al*., 2014; Woessner & Goodenough, 1994;). Whether their glycosylation, showing similarities to land plant extensins (Bollig *et al*., 2007), is performed by homologs of land plant enzymes from the GT95, GT77 and GT47 families is questionable by looking at the work of Barolo *et al*. (2020) who detected few, if any, homologs in microalgal genomes. (iii) The complex interplay of three different P4Hs (AtP4H2, AtP4H5 and AtP4H13) was shown in *A. thaliana* (Velasquez *et al*., 2015). The formation of homodimers of AtP4H5 or heterodimers of AtP4H5 with either AtP4H2 or AtP4H13 is required for correct peptidyl-proline hydroxylation. This is the case for at least extensin, but cannot be ruled out for other HRGPs. From these three P4Hs, GBG85197 showed most similarity with AtP4H13 in the βII-βIII loop thus hinting potential inactivity in a monomeric state. All these three possible explanations are speculative but highlight the general aim that presence or absence of P4H homologs should not be seen equivalent to presence or absence of AGPs or extensins. The latter has to be individually shown by a combination with analytical methods as shown in this study.

## Conclusion

Although presence of pectic polysaccharides is undoubtful and especially the walls of *C. aspera* contain huge amounts of these polysaccharides possibly involved in salt tolerance of this species, further investigations have to follow to elucidate the exact structural composition and to clarify, whether only homogalacturonan or also RG-I or even RG-II have evolutionary roots in algae. The most prominent unusual structural features detected in the aqueous extracts of the investigated Charales were 1,4-linked rhamnans and 1,3-, 1,6- and 1,3,6-linked partially methylated galactans. Obviously these galactans were not part of AGPs. Although one possible functional sequence of P4H was detected in the genome of *Chara braunii*, no Hyp (necessary for linkage of the galactan to the protein moiety in AGPs) was detected in any of the investigated species. The fact, that antibodies directed against AGP epitopes showed cross reactivities with the unusual methylated galactans is a reminder that results only based on antibodies are not always reliable. Our work reveals strong differences of the cell walls of members of the Charophyceae compared to seed plants and even to the other members of the higher-branching ZCC-grade of streptophyte algae, namely the Coleochaetophyceae and the Zygnematophyceae. More detailed investigations of the cell walls of the different streptophyte algal taxa are necessary to understand how these algae met the challenge of cell wall adaptation necessary for life on land.

## AUTHORS CONTRIBUTIONS

BC and LP planned and designed the research. HS collected and identified the plant material. FE, KM, LP and JU performed the extractions as well as carbohydrate and ELISA experiments. JBJZ performed preliminary analytical work on *Chara* species from Austria. LP performed bioinformatic searches. LP and KM created all main text figures. All authors analysed the data (BC, FE, KM, LP: carbohydrate analysis; LP bioinformatics search for prolyl 4-hydroxylase). BC, KM and LP wrote the draft manuscript; all authors revised the manuscript, read and approved the final manuscript.

## ACKNOWLEDGEMENTS

The authors thank Prof. Dr. Michael Schagerl and Barbara Mähnert from the Department for Limnology and Oceanography of the University of Wien for collection of *Chara* species used in preliminary experiments. Funding: LP and BC (project-number 440046237) are grateful for funding within the framework of MAdLand (http://madland.science), priority programme 2237 of the German Research Foundation (DFG). HS also thanks the DFG for funding (project SCHU 983/23-1).

## DATA AVAILABILITY STATEMENT

All relevant data can be found within the manuscript and its supporting materials.

## SUPPORTING INFORMATION

Additional supporting information may be found online in the Supporting Information section at the end of the article:

**TABLE S1.** Antibodies tested for binding to *Chara* spp. cell wall fractions AE, AmOx or KOH.

**TABLE S2.** Yields of the different extracts from different *Chara* species in % of dry plant material (w w^-1^).

**TABLE S3a.** Neutral monosaccharide composition of water-soluble polysaccharides (AE) from different *Chara* species and *Nitellopsis obtusa* (Pfeifer *>et al.*, 2022) in % (mol mol^-1^ tr: trace value < 1%).

**TABLE S3b.** Neutral monosaccharide composition of dialyzed water-soluble polysaccharides, pre-treated with amylase and pectinase (AE_AP), from different *Chara* species and *Nitellopsis obtusa* in % (mol mol^-1^ tr: trace value < 1%).

**TABLE S4a.** Detailed comparison of the oxoglutarate-binding region of putative *Chara braunii* P4Hs. Amino acid positions refer to P4H from *Chlamydomonas reinhardtii* according to Koski *et al*. (2007).

**TABLE S4b.** Detailed comparison of the iron-binding region of putative *Chara braunii* P4Hs. Amino acid positions refer to P4H from *Chlamydomonas reinhardtii* according to Koski *et al*. (2007).

## Notes

### Competing Interest Statement

The authors have declared no competing interest.

