## Supporting Information for "The cell walls of different *Chara* species (Charophyceae) are characterized by branched galactans rich in 3-*O*-methylgalactose and absence of arabinogalactan-proteins"

[+]Authors contributed equally to the publication.

*Author for correspondence:

**TABLE S1** Antibodies tested for binding to *Chara* spp. cell wall fractions AE, AmOx or KOH.

| Antibody | Epitope | Key References |
|---|---|---|
| INRA-RU2 | [(4)-α-D-GalA-(1,2)-α-L-Rha]$_4$ of RG-I | Ralet *et al*. (2010) |
| JIM13 | AGP glycan,<br>e.g. β-D-GlcA*p*-(1→3)-α-D-GalA*p*-(1→2)-α-L-Rha | Pfeifer *et al*. (2022);<br>Yates *et al*. (1996) |
| KM1 | (1→6)-β-D-Gal*p* units in AGs type II | Classen *et al*. (2004);<br>Ruprecht *et al*. (2017) |
| LM2 | (1→6)-β-D-Gal*p* units with terminal ß-D-GlcA*p* in AGP | Ruprecht *et al*. (2017);<br>Smallwood *et al*. (1996); |
| LM6 | (1→5)-α-L-Ara*f* oligomers in arabinan or AGP | Verhertbruggen *et al*. (2009a) |
| LM10 | Non-reducing end of (1→4)-β-D-xylan | McCartney *et al*. (2005);<br>Ruprecht *et al*. (2017) |
| LM15 | XXXG-motif of xyloglucan | Marcus *et al*. (2008);<br>Pedersen et al. (2012) |
| LM19 | Unesterified HG | Verhertbruggen *et al*. (2009b) |

**TABLE S2** Yields of the different extracts from different *Chara* species in % of dry plant material (w w$^{-1}$).

| | *C. aspera* | *C. globularis* | *C. subspinosa* | *C. tomentosa* |
|---|---|---|---|---|
| AE | 2.5 | 0.6 | 1.6 | 1.3 |
| $(NH_4)_2C_2O_4$ | 21.2 | 2.3 | 5.9 | 5.0 |
| HCl | 1.9 | 0.3 | 0.2 | 0.7 |
| $Na_2CO_3$ | 2.9 | 0.7 | 1.2 | 1.0 |
| KOH | 6.1 | 6.3 | 3.5 | 2.8 |

**TABLE S3a** Neutral monosaccharide composition of water-soluble polysaccharides (AE) from different *Chara* species and *Nitellopsis obtusa* (Pfeifer *et al*., 2022) in % (mol $mol^{-1}$; tr: trace value < 1%).

| Neutral mono-saccharide | *C. aspera* AE n=3 | *C. globularis* AE n=3 | *C. subspinosa* AE n=3 | *C. tomentosa* AE n=3 | *N. obtusa* AE n=3 |
|---|---|---|---|---|---|
| 3-*O*-MeRha | tr | - | - | - | - |
| Rha | 11.8 ± 0.2 | 10.7 ± 0.9 | 11.9 ± 0.4 | 10.3 ± 0.8 | 18.0 ± 0.4 |
| 3-*O*-MeFuc | tr | tr | 1.9 ± 0.1 | 1.1 ± 0.2 | - |
| Fuc | 7.0 ± 0.7 | 12.1 ± 2.4 | 7.2 ± 0.1 | 11.0 ± 1.1 | 5.8 ± 0.0 |
| Rib | tr | 1.4 ± 0.2 | tr | tr | - |
| Ara | 3.0 ± 0.1 | 6.9 ± 1.9 | 3.4 ± 0.1 | 6.4 ± 2.8 | 20.8 ± 0.7 |
| Xyl | 6.1 ± 0.3 | 9.6 ± 1.3 | 16.4 ± 0.5 | 12.8 ± 1.5 | 3.7 ± 0.5 |
| Man | 8.7 ± 0.4 | 9.1 ± 0.3 | 7.2 ± 0.4 | 12.3 ± 5.0 | 7.3 ± 0.4 |
| 3-*O*-MeGal | 18.8 ± 0.3 | 5.8 ± 1.6 | 7.4 ± 0.2 | 3.7 ± 0.7 | - |
| Gal | 13.7 ± 0.4 | 22.9 ± 2.8 | 13.5 ± 0.3 | 12.7 ± 1.6 | 18.6 ± 0.2 |
| Glc | 30.9 ± 1.4 | 21.5 ± 1.6 | 31.1 ± 0.6 | 29.7 ± 2.4 | 25.8 ± 0.9 |

**TABLE S3b** Neutral monosaccharide composition of dialyzed water-soluble polysaccharides, pre-treated with amylase and pectinase (AE_AP), from different *Chara* species and *Nitellopsis obtusa* in % (mol $mol^{-1}$; tr: trace value < 1%).

| Neutral mono-saccharide | *C. aspera* AE_AP n=3 | *C. globularis* AE_AP n=3 | *C. subspinosa* AE_AP n=3 | *C. tomentosa* AE_AP n=3 | *N. obtusa* AE_AP n=3 |
|---|---|---|---|---|---|
| Rha | 15.5 ± 0.2 | 12.2 ± 0.2 | 14.5 ± 0.2 | 10.0 ± 0.1 | 21.8 ± 0.3 |
| 3-*O*-MeFuc | tr | tr | 1.8 ± 0.1 | 1.0 ± 0.1 | - |
| Fuc | 9.7 ± 0.2 | 16.5 ± 0.3 | 9.8 ± 0.2 | 14.3 ± 0.1 | 7.4 ± 0.3 |
| Ara | 2.4 ± 0.2 | 4.0 ± 0.1 | 2.5 ± 0.0 | 2.2 ± 0.0 | 12.5 ± 0.1 |
| Xyl | 7.5 ± 0.4 | 9.5 ± 0.4 | 10.1 ± 0.2 | 14.3 ± 0.5 | 5.2 ± 0.2 |
| Man | 8.5 ± 0.2 | 9.0 ± 0.2 | 10.1 ± 0.1 | 12.0 ± 0.3 | 8.0 ± 0.2 |
| 3-*O*-MeGal | 19.3 ± 0.2 | 6.6 ± 0.3 | 9.9 ± 0.2 | 4.2 ± 0.0 | 3.4 ± 0.1 |
| Gal | 16.8 ± 0.2 | 22.4 ± 0.2 | 20.3 ± 0.2 | 14.5 ± 0.1 | 27.2 ± 0.4 |
| Glc | 20.3 ± 0.5 | 19.8 ± 0.4 | 21.0 ± 0.1 | 27.5 ± 0.4 | 14.5 ± 0.7 |

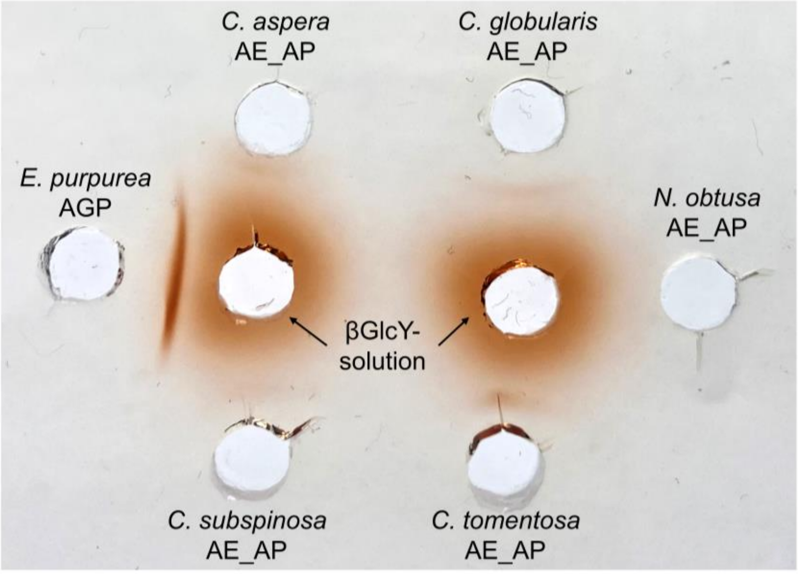


**FIGURE S1** Gel diffusion assay with aqueous fractions after enzymatic treatment from *C. globularis, C. subspinosa*, *C. tomentosa* and *C. aspera* and *N. obtusa* (100 mg $mL^{-1}$) and βGlcY (1 mg $mL^{-1}$). The red precipitation line indicates presence of AGPs. As positive control, AGP from *Echinacea purpurea* (10 mg $mL^{-1}$) was used.

**TABLE S4a** Detailed comparison of the oxoglutarate-binding region of putative *Chara braunii* P4Hs. Amino acid positions refer to P4H from *Chlamydomonas reinhardtii* according to Koski *et al*. (2007).

| | $Tyr_{134}$ | $Thr_{178}$ | $Lys_{237}$ | $Ser_{239}$ | $Thr_{241}$ | $Trp_{243}$ |
|---|---|---|---|---|---|---|
| GBG60506 | ✓ | × (Gln) | ✓ | × (Lys) | × (Lys) | × (Leu) |
| GBG60917 | × | × | × | × | × | × |
| GBG70085 | ✓ | ✓ | ✓ | ✓ | ✓ | ✓ |
| GBG70087 | ✓ | × (Met) | ✓ | × (Lys) | × (Lys) | × (Leu) |
| GBG72676 | × | ✓ | ✓ | ✓ | ✓ | ✓ |
| GBG72677 | × | ✓ | ✓ | ✓ | ✓ | ✓ |
| GBG85197 | ✓ | ✓ | ✓ | ✓ (Val) | ✓ | ✓ |

**TABLE S4b** Detailed comparison of the iron-binding region of putative *Chara braunii* P4Hs. Amino acid positions refer to P4H from *Chlamydomonas reinhardtii* according to Koski *et al*. (2007).

| | $His_{143}$ | $Asp_{145}$ | $His_{227}$ |
|---|---|---|---|
| GBG60506 | ✓ | ✓ | ✓ |
| GBG60917 | × | × | × |
| GBG70085 | ✓ | ✓ | ✓ |
| GBG70087 | × (Met) | × | × |
| GBG72676 | × | × | ✓ |
| GBG72677 | × | × | ✓ |
| GBG85197 | ✓ | ✓ | ✓ |
